# Strengthening Antisense Oligonucleotide-Mediated Anti-Tumor Immunity via Metal-Organic Framework Nanoparticles

**DOI:** 10.1101/2025.03.28.645811

**Authors:** Julia A. Nowak, Ezra Cho, Meredith A. Davis, Siyi Zheng, Lauren Bell, Fanrui Sha, Julian S. Magdalenski, Omar K. Farha, Michelle H. Teplensky

**Affiliations:** Department of Biomedical Engineering, Boston University; Division of Materials Science and Engineering, Boston University; Department of Chemistry, Northwestern University

## Abstract

Overexpression of checkpoint proteins, such as programmed death ligand one (PD-L1), prevents immune recognition and enables cancer growth. Current monoclonal antibodies that block PD-L1 tend to be fragile, unable to penetrate tumors, and target cancer at later stages, thus leading to inconsistent patient outcomes. Antisense oligonucleotides (ASOs) provide an alternative to decrease PD-L1 expression, but require frequent high dosing due to fast degradation, rapid clearance, and poor cell uptake. To overcome these issues, we harnessed biocompatible metal-organic framework (MOF) nanoparticles, porous nanomaterials comprising metal nodes and organic linkers, to deliver ASOs. Encapsulating ASOs into MOFs enhances their stability and protection during intracellular delivery, leading to reduced PD-L1 expression and downstream immune recognition. Herein, we synthesized three distinct PD-L1-specific ASOs and loaded them individually into zirconium-based nano-sized NU-1000 MOFs, averaging ∼80% encapsulation efficiency. Release of encapsulated ASOs was sustained up to 7 days *ex cellulo*. MOF encapsulation increased ASO potency and reduced PD-L1 expression *∼*3-fold and 2-fold in triple negative breast cancer EMT6 and melanoma B16-F10 cells, respectively. We evaluated the impact of MOF-delivered ASOs on PD-L1-expressing immune cells, where we observed *ca.* 12-fold increases in dendritic cell co-stimulatory marker expression, and amplified T cell activation and proliferation compared to untreated cells (4-fold and 10-fold, respectively). Notably, these changes drove a 3-fold increase in tumor caspase-3 expression, a key mediator for apoptosis. This research highlights how MOFs can be harnessed to bypass ASO limitations without requiring sequence modifications, and offers a broadly applicable platform for improved oligonucleotide delivery for various genes of interest.

## Introduction

Overexpression of checkpoint proteins, such as programmed death ligand one (PD-L1), on cancer cells enables unregulated tumor growth and immune evasion.^1–3^ Decreasing PD-L1/Programmed death protein 1 (PD-1) recognition is a clinically employed strategy to boost immune recognition of cancer and subsequently activate an antitumor response.^1,4^ Indeed, existing immunotherapies such as Tecentriq, Bavencio, and Imfinzi, are monoclonal antibodies that target and block PD-L1 to improve immune recognition of cancer.^5^ While successful against some cancers (*e.g.,* merkel cell carcinoma), these therapies are generally limited by their large size, instability, and poor tumor penetration.^5–9^ Such disadvantages can ultimately lead to poor and inconsistent patient outcomes across various cancers, such as in melanoma or triple negative breast cancer, which do exhibit elevated PD-L1 as a therapeutic target.^7–10^ Therefore, there is a need for alternative therapies that target PD-L1 expression on cancer cells and promote increased immune activity.

One such alternative is antisense oligonucleotides (ASOs).^11,12^ ASOs can selectively target and inhibit protein expression of key genes of interest, such as PD-L1.^11–14^ Their specific binding to gene targets decreases the subsequent expression of proteins; this is an advantage over antibodies, which bind to already expressed PD-L1 proteins and solely block interactions with their receptors.^15–17^ In contrast, ASOs can reduce the probability of PD-L1/PD-1 interactions by decreasing surface expression of the PD-L1 protein. However, ASOs are limited due to rapid clearance, fast degradation, and poor cellular uptake.^11,18^ Chemical modifications to an ASO can increase stability and nuclease resistance, but extensive modifications can decrease target RNA affinity and adversely affect potency.^19–21^ Alternatives to increase free ASO uptake include conjugation of ASOs to lipids or peptides, or the use of nanoparticle delivery systems.^11,18,22^ However, such changes risk alteration to ASO binding affinity, or require local delivery or functionalization to access and target cells beyond the liver or reticuloendothelial system.^11,18^ As such, we explored a broadly translatable delivery method that offers stability, high uptake, and can achieve high ASO loading capacities to reduce the required dose and frequency of dosage. Herein, we employed metal-organic framework (MOF) nanoparticles to elevate ASO protection, uptake, and delivery into cells.

MOFs are highly porous nanoparticles that allow for tuned release of encapsulated biological cargo at significantly higher loadings (up to 40 wt.%) than common delivery systems.^23–27^ MOFs comprise metal nodes coordinated to organic linkers, and possess high modularity that can be harnessed to tailor drug delivery and uptake into various cell types.^27–30^ In this work, we used the biocompatible zirconium-based MOF, NU-1000, whose nano-scale size (100 – 200 nm) and channel-like structure can successfully encapsulate biological cargo. Thus, its high surface area, porosity, and biocompatibility is well-suited for sustained ASO delivery.

Herein, we illustrate how MOF-mediated delivery of PD-L1-targeting ASOs successfully decreases PD-L1 expression and stimulates antitumor immunity. We selected three unique ASO sequences to target the PD-L1 gene and successfully encapsulated each within nano-sized NU-1000 particles. We achieved high encapsulation efficiencies and overall loadings, *ca.* 80% and 9 nmol ASO per mg of MOF, respectively. We sustained release of ASO from NU-1000 for over 7 days, characterized by an initial 12 hours of release followed by a slower, detained release profile. MOF delivery improves ASO cell uptake >3-fold and decreases PD-L1 surface expression on cancer cells compared to free ASO. Notably, ASO encapsulated within MOF (termed ASO@MOF) lowers surface PD-L1 levels to those at or below unstimulated melanoma (B16-F10) and triple negative breast (EMT6) carcinoma cells. Furthermore, ASO@MOF propagated potent anti-tumor immunity. Namely, ASO@MOF, but not free ASO or unloaded MOF, stimulated dendritic cell co-stimulatory marker expression by *ca.* 2 to 10-fold, and increased T cell activation marker expression and subsequent proliferation by *ca.* 4 and 10-fold, respectively. Overall, this work highlights how MOF-mediated delivery of PD-L1-targeting ASOs: 1) protects the oligonucleotides from degradation and elongates their release profile, 2) elevates ASO uptake into relevant cells, and 3) enables increased tumor susceptibility to immune attack due to downregulated PD-L1 expression. These results have the potential to positively impact drug delivery and elicit greater anti-tumor immunity in PD-L1-overexpressing cancers beyond melanoma or triple negative breast cancer, and can also be applied to other genes of interest that are overexpressed across cancer (*e.g.,* CD47^29^ or EGFR^1,31^).

## Results

### Design and Encapsulation of PD-L1-Targeting ASOs

We designed ASOs targeting the gene of interest, PD-L1, by evaluating sequences from prior literature to ensure gene specificity.^13,14^ We chose sequences with lengths of *ca.* 20 base pairs, termed ASO A, ASO B, and ASO C (Table S1).^13,14^ Each sequence is reported to target PD-L1, with ASO B additionally incorporating CG motif repeats capable of activating toll-like receptor 9 (TLR9), which triggers an innate immune response.^13,14^ After standard solid-phase phosphoramidite synthesis and purification of the ASOs, we characterized the molecular weight with matrix-assisted laser desorption/ionization time-of-flight (MALDI-TOF) (Table S1).

We employed 100-200 nm NU-1000 MOF nanoparticles (Fig. S1). This mesoporous framework, which contains a large 3.3 nm pore aperture^32^ than many conventionally used MOFs (*e.g.,* UiO-66^25^, MIL-88 (Fe)^33^), can internalize greater amounts of biomacromolecular cargo post-synthesis.^33^ For this reason, consistent with prior literature illustrating the feasible internalization of oligonucleotides within a pore of this diameter,^34^ we hypothesized that the NU-1000 MOF could encapsulate high amounts of ASO. We encapsulated ASO A, ASO B, or ASO C into NU-1000, and observed *ca*. 80% efficiency across all sequences (Fig. 1A). The loading was equivalent to *ca.* 9 nmol/mg MOF, or 20 wt.%, which is a higher ratio than typical loadings in conventionally-employed lipid or polymer-based nanoparticles (*i.e.,* 5 wt.%^33^). To evaluate the timeframe of encapsulated ASO release from the MOFs, we synthesized Cy3 dye-labelled ASO A, and measured release from MOF in a phosphate-buffered saline (PBS) solution over time (Fig. 1B). We observed an initial rapid release for the first 12 hours, where *ca.* 50% of the loaded cargo was released into the solution. This was followed by a sustained release of the remaining encapsulated Cy3 dye-labelled ASO A over the course of 7 days. We suggest that the initial burst release consists of cargo closest to the pore channel opening that diffuses out of the MOF channels. Then, as previously reported,^34^ gradual MOF disassembly occurs through phosphate-mediated linker displacement, facilitating the release of further internally-loaded cargo. The measured elongated ASO release profile is advantageous for multiple reasons. It: 1) ensures a sustained concentration of desired drug and 2) lowers the need for multiple drug administrations, as a single MOF nanoparticle can release ASO for up to 7 days.

**Figure 1.**
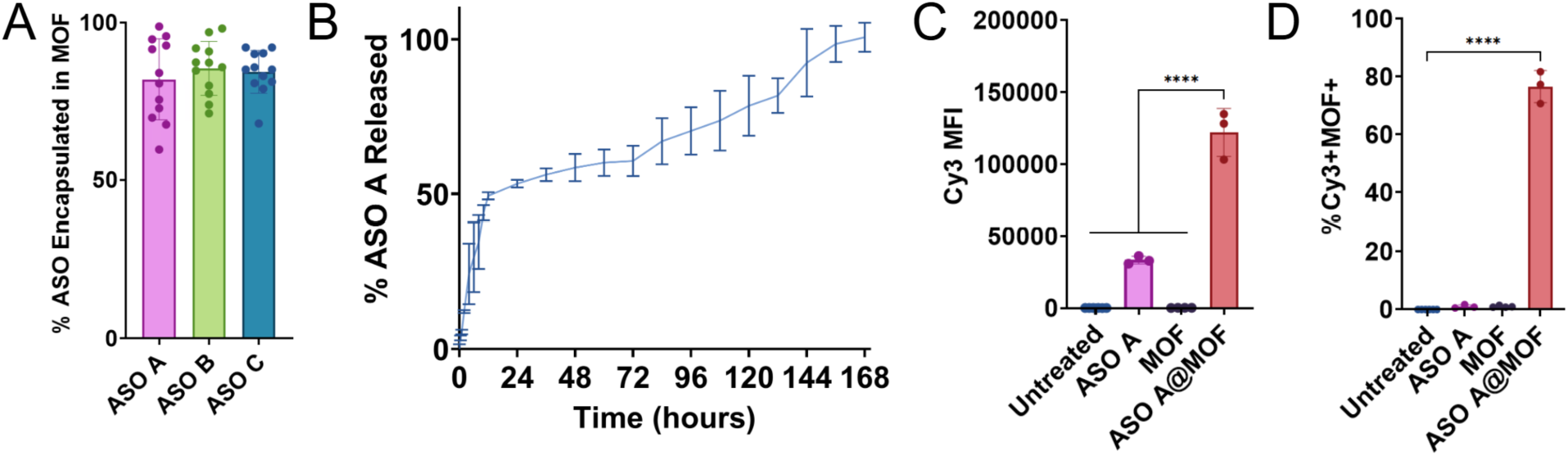
Successful encapsulation, release, and cellular uptake of ASO@MOF. (**A**) Consistent loading of diverse PD-L1 specific ASO sequences in NU-1000 MOF. N= 12 (**B**) Sustained release of Cy3 dye-labelled ASO A over 7 days. (**C**) Cy3 dye-labelled ASO A@MOF exhibits *ca.* 3-fold increased cellular uptake into B16-F10 cells and (**D**) entry into >75% of cell population (n = 3 - 4). Analysis was performed using an ordinary one-way ANOVA, followed by a Tukey’s multiple comparisons test. Data show mean ± s.d. ****p<0.0001.

### Elevated ASO Intracellular Uptake via MOF Nanoparticle Delivery

Given an understanding of how the ASO@NU-1000 nanoparticles release their cargo, we next sought to evaluate how this timescale compares with the rate of intracellular uptake. When incubated with B16-F10 cells, the MOF increased encapsulated ASO A uptake by >3-fold compared to free ASO A in solution (Fig. 1C). ASO A encapsulated in MOF (ASO A@MOF) entered more than 75% of the total B16-F10 cell population (Fig. 1D). We also measured MOF uptake into the cells and observed MOF entry to be cell-type independent. Similarly high uptake for MOF-encapsulated ASOs was measured across melanoma B16-F10 and breast cancer EMT6 cells as well as in immune cells such as bone marrow-derived dendritic cells (BMDCs) (∼60% or more of the cell population) (Fig. S2).

### MOF-elevated ASO Uptake Decreases PD-L1 Surface Expression

After evaluating the ability of MOF encapsulation to elevate ASO uptake across various cell types, we assessed how ASO@MOF delivery strengthened ASO potency, specifically the downregulation of PD-L1 surface expression. Increased PD-L1 surface expression on B16-F10 and EMT6 cells was induced using interferon gamma (IFNγ) prior to ASO@MOF treatment. The ASO sequences chosen were not designed to be cell type-specific, and therefore, we hypothesized that we would be able to reduce PD-L1 expression across multiple cancer types. In B16-F10 cells, ASO A@MOF treatment and ASO B@MOF treatment led to *ca.* 2.5-fold decreased PD-L1 surface expression, when compared to the IFNγ-induced cell control (Stimulated/Untreated), even bringing PD-L1 median fluorescence intensity (MFI) down to unstimulated levels (∼2000) (Fig. 2A). ASO C@MOF treatment had no significant effect on PD-L1 expression. Treatment with free ASO, regardless of the specific sequence, did not decrease PD-L1 expression in B16-F10 or EMT6 cells, and, in contrast, elevated PD-L1 expression in some cases (Fig. 2A, B). We suggest this is due to inefficient uptake of ASO when delivered freely and concurrent increased expression of the PD-L1 receptor by the cancer cells as a defense mechanism against treatment modulation. In EMT6 cells, ASO A@MOF treatment led to a *ca.* 2-fold decrease in PD-L1, restoring it completely to unstimulated levels (Fig. 2B). The impact from ASO B@MOF and ASO C@MOF treatment was significantly greater, causing a *ca.* 8-fold decrease in PD-L1 surface expression compared to the IFNγ-induced cell control group (Stimulated/Untreated), and even decreasing PD-L1 expression to 4-fold below that of the unstimulated control group (Untreated) (Fig. 2B). Despite the broad PD-L1-targeting capabilities of the ASO sequences chosen, we suggest there are differences in how they are intracellularly trafficked and metabolized, which is what we attribute to the differences in sequence knockdown potency measured between B16-F10 and EMT6 cell lines. Overall, while different ASO@MOF treatments altered the extent of reduced PD-L1 surface expression across cell types, the significant decrease in surface PD-L1 compared to free ASO, unloaded MOF vehicle, and untreated cells highlights how ASO@MOF is effective as a platform to deliver fragile yet potent ASOs to target a robust cancer immune suppression mechanism.

**Figure 2.**
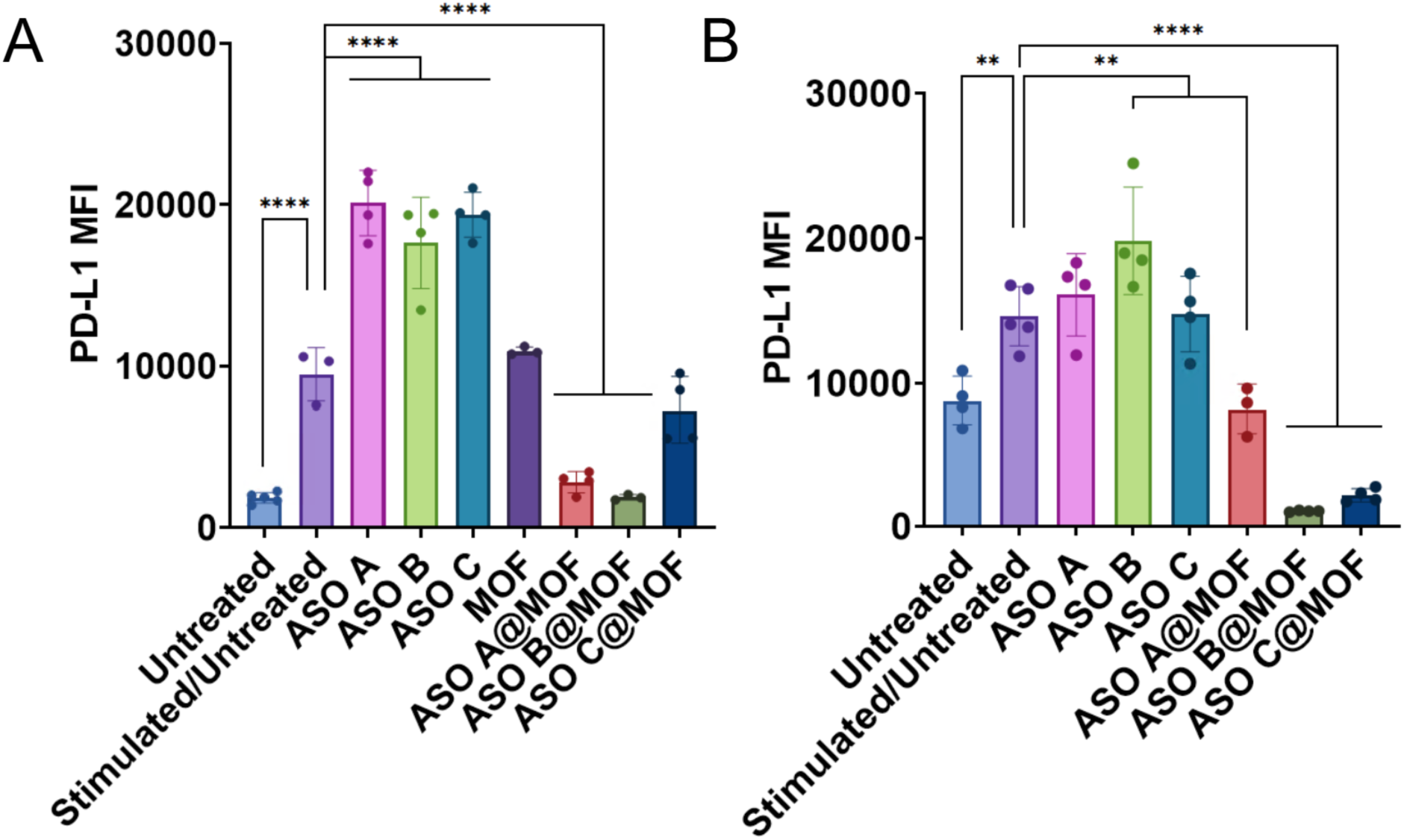
ASO@MOF treatment of (**A**) B16-F10 melanoma cells and (**B**) EMT6 triple negative breast cancer cells successfully downregulates PD-L1 expression compared to untreated control stimulated to express elevated PD-L1 (n = 3 – 4). Free ASO or unloaded MOF exhibit increased or no change in PD-L1 expression. Analysis was performed using an ordinary one-way ANOVA, followed by a Tukey’s multiple comparisons test. Data show mean ± s.d. **p<0.01; ****p<0.0001.

### ASO@MOF Increases Immune Cell Activation and Proliferation

To understand how ASO@MOF impacts other PD-L1-expressing immune cell populations alongside its desired effects on cancer cells, we analyzed surface presentation of immune activation markers and proliferation after treatment. We observed minimal innate stimulatory effects of the free ASOs compared to immunostimulant Class B CpG oligonucleotide 1826 using a macrophage reporter cell line (RAW-Blue) (Fig. S3). We next evaluated how this immunostimulation changed when these ASOs were delivered via MOFs.

To evaluate the effects of ASO@MOF on mortal immune cells, we treated and assessed responses in BMDCs, whose PD-L1 presentation can dampen anti-tumor immunity.^35^ Through MOF-mediated delivery, the various ASO sequences increased BMDC activation, as measured through flow cytometry analysis of CD80 and CD86 co-stimulatory marker expression (Fig. 3A, B). CD80 and CD86 exhibited *ca.* 2-fold and *ca.* 10-fold increases in MFI, respectively, compared to untreated and free ASO-treated BMDCs (Fig. 3A, B). ASO A@MOF induced the highest increase in both CD80 and CD86 expression, which does correspond to the minor increase measured in the RAW-Blue reporter cell line with free ASO A at the highest tested concentration (2000 μM) (Fig. S3). However, the significant increases in co-stimulatory marker expression indicate that MOF-mediated delivery provides an additional immune benefit that is not present with free ASO. We hypothesize that this stimulatory immune response from ASO@MOF treatment will further promote the ability to elicit an anti-tumor immune response.

**Figure 3.**
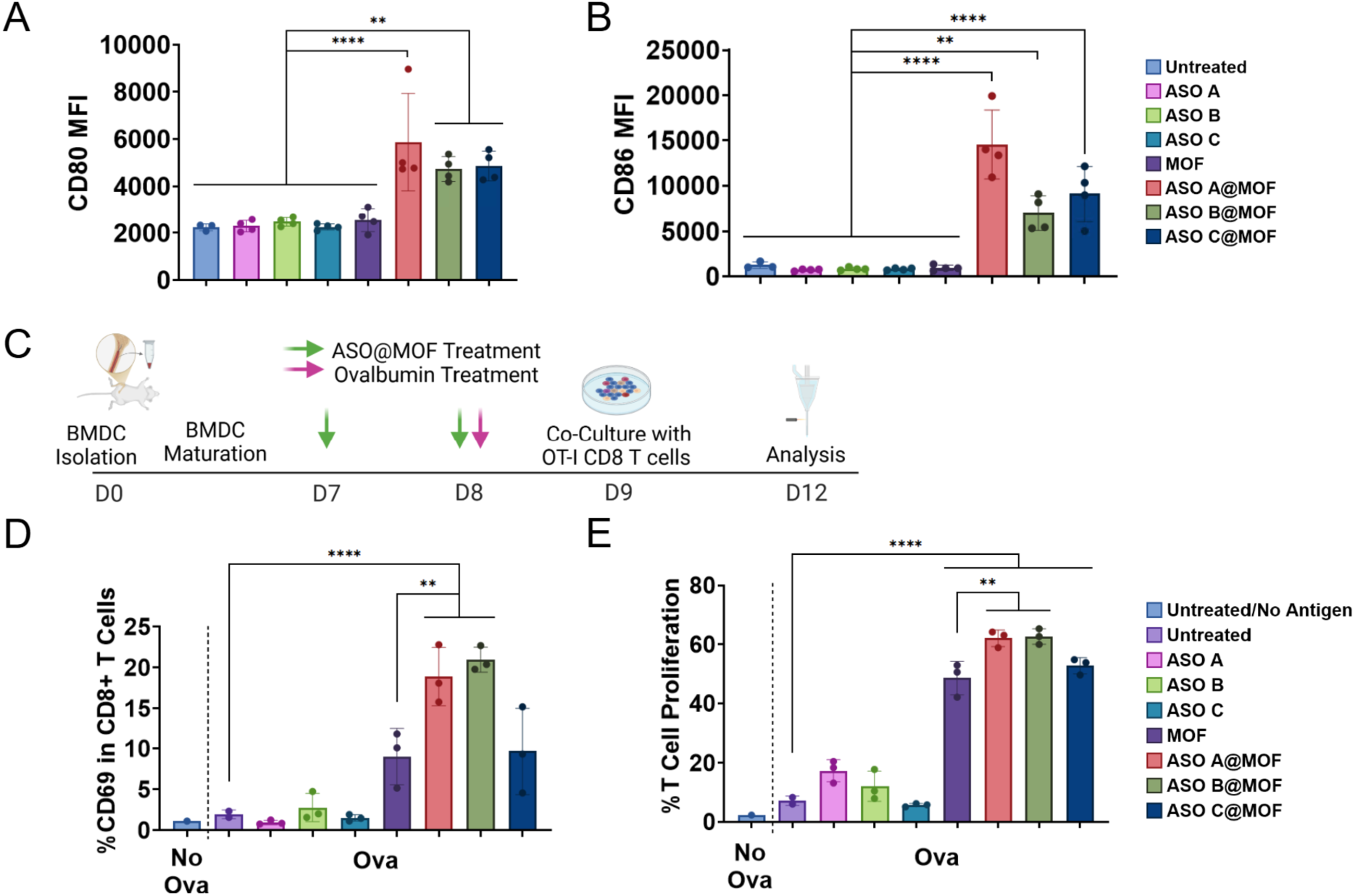
ASO@MOF treatment increases immune cell activation and amplifies subsequent T cell proliferation. BMDCs treated with ASO@MOF exhibit increased co-stimulatory (**A**) CD80 and (**B**) CD86 marker expression compared to untreated, free ASO, and unloaded MOF (n = 4). (**C**) Experimental schematic showing that upon co-culture with OT-I CD8^+^ T cells, (**D**) further increases in early T cell activation marker CD69 and (**E**) T cell proliferation occur for ASO@MOF treated BMDCs (n = 2 – 3). Analysis was performed using an ordinary one-way ANOVA, followed by a Tukey’s multiple comparisons test. Data show mean ± s.d. **p<0.01; ****p<0.0001.

To understand the downstream effects on the immune environment propagated through ASO@MOF treatment due to altered PD-L1 expression, we analyzed subsequent T cell activation and proliferation (Fig. 3C). After treating BMDCs with various free ASO or ASO@MOF alongside model protein antigen ovalbumin (OVA), we co-cultured the treated BMDCs with OT-I CD8^+^ T cells, which are specific to the OVA1 (SIINFEKL) epitope within the OVA protein. As such, measured changes in T cell activation and proliferation are indicative of changes in the interaction between CD8^+^ T cells and treated BMDCs that are importantly affected by PD-L1 expression (Gating strategy, Fig. S4). We observed a *ca.* 4-fold increase in early activation marker CD69 expression when BMDCs were treated with MOF (Fig. 3D). ASO A@MOF and ASO B@MOF significantly elevated the extent of CD69 expression amongst CD8^+^ T cells (*ca.* 20% of the CD8^+^ T cell population expressed CD69) compared to groups where BMDCs were treated with unloaded MOF (*ca.* 8% of CD8^+^ were CD69^+^) or free ASO (*ca.* 2-3% of CD8^+^ were CD69^+^). This signifies that MOF-mediated delivery of ASO A and ASO B generates a BMDC response that better activates CD8^+^ T cells. Moreover, the elevated T cell activation plays an essential role in anti-tumor immunity through robust proliferation of an antigen-specific CD8^+^ T cell population. ASO@MOF treatment positively impacted BMDC-antigen-specific T cell interactions, which increased antigen-specific CD8^+^ T cell proliferation (*ca.* 60-65% proliferation) (Fig. 3E). The presence of MOF in the cell co-culture increased CD8^+^ T cell proliferation to *ca.* 45%, possibly attributed to free OVA in the co-culture solution being encapsulated into unloaded MOF and subsequently internalized by BMDCs (Fig. 3E). Importantly, however, ASO A@MOF and ASO B@MOF further elevated CD8^+^ T cell proliferation by an additional *ca.* 20%. This illustrates how MOF encapsulation of PD-L1-specific ASOs can increase ASO potency of gene knockdown in a cancer population while also improving anti-tumor immunity among immune cell populations.

### ASO@MOF Treatment Increases Cancer Cell Apoptosis

Ultimately, we explored how ASO@MOF treatment affects the resulting apoptosis of cancer cells. We hypothesized that downstream impact of reduced PD-L1 expression in addition to enhanced immune activation would improve immune-mediated tumor cell death. B16-F10 cells were stimulated with IFNγ to induce PD-L1 expression and treated with various ASO@MOFs to knockdown that expression. Treated cells were then co-cultured with Pmel-1 splenocytes, which have T cells specific for the gp-100 epitope expressed by B16-F10 cells, and tumor cells were analyzed for expression of Caspase-3, a key mediator in cell apoptosis (Fig. 4A). Notably, compared to the untreated PD-L1-stimulated B16-F10 cells (Stimulated/Untreated), we measured a *ca.* 3-fold increase in Caspase-3 expression after treatment with ASO A@MOF, ASO B@MOF, or ASO C@MOF (Fig. 4B). Treatment with free ASO or unloaded MOF did not significantly increase Caspase-3 expression (Fig. 4B). Furthermore, the percentage of B16-F10 cells expressing active Caspase-3 increased to >30% of the total population in ASO@MOF-treated groups, a statistically significant increase compared to the untreated baseline of *ca.* 15% active Caspase-3 (Fig. 4C). There were no significant differences measured in the percentage of cells with active Caspase-3 treated with free ASO or unloaded MOF compared to untreated cells (Stimulated/Untreated). Holistically, these data highlight that irrespective of PD-L1 sequence, MOF encapsulation of PD-L1-targeting ASO meaningfully downregulates PD-L1 expression, increases immune stimulation, and propagates beneficial anti-tumor interactions that drive immune cell-mediated apoptosis of B16-F10 cancer cells.

**Figure 4.**
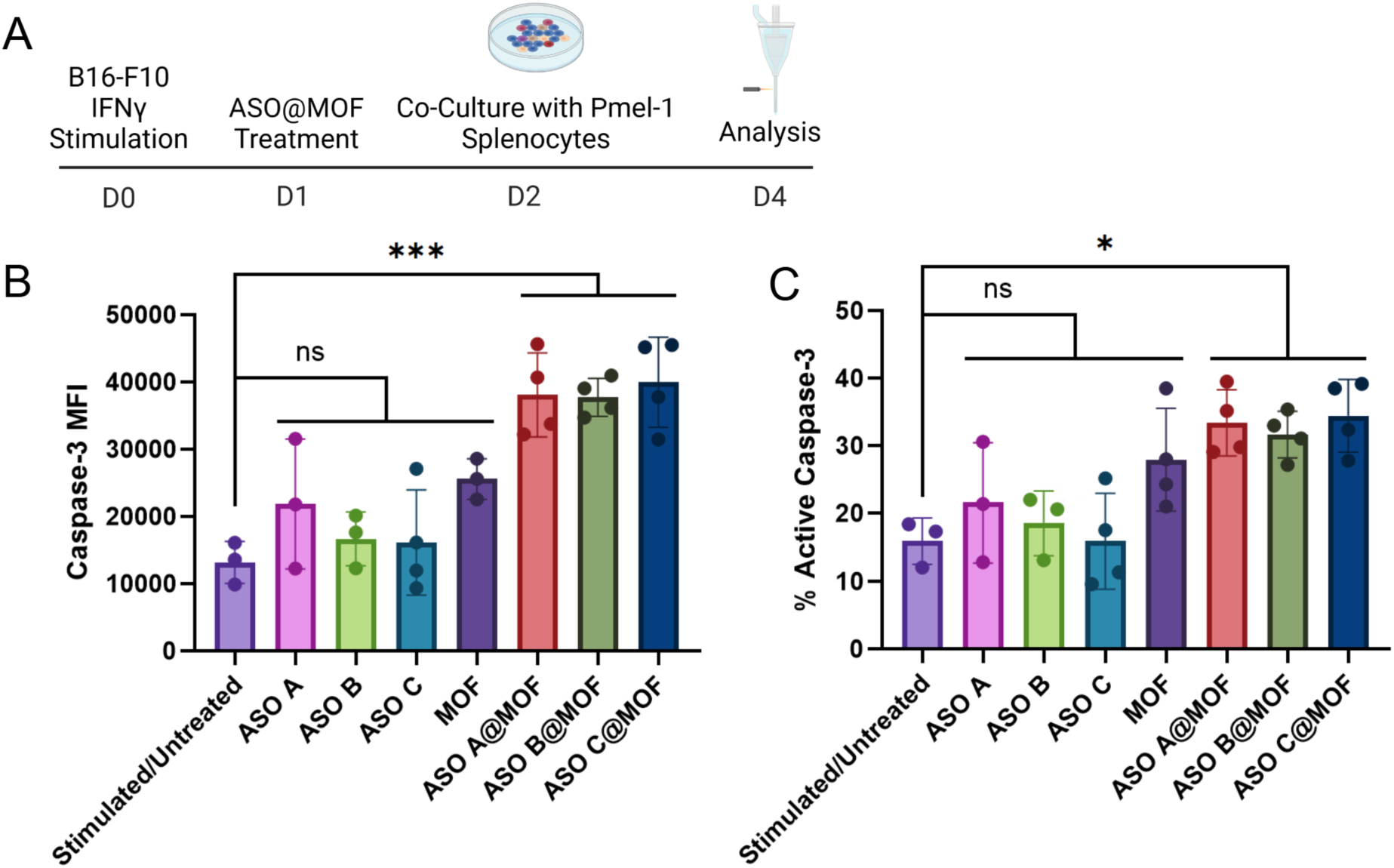
ASO@MOF increases cancer cell apoptosis. (**A**) Experimental treatment schedule. (**B**) Treatment of B16-F10 cells with ASO@MOF increases early cell death Caspase-3 marker expression in tumor cells after co-culture with Pmel-1 splenocytes, and (**C**) percentage of tumor cells expressing active Caspase-3 in comparison to untreated control (n = 3 – 4). Free ASO or unloaded MOF showed no significance in comparison to untreated control for both (**B**) Caspase-3 expression and (**C**) percent cells with active Caspase-3. Analysis was performed using an ordinary one-way ANOVA, followed by a Tukey’s multiple comparisons test. Data show mean ± s.d. *p<0.05; ***p<0.001; ns=nonsignificant.

## Discussion

This work explores the implementation of MOF nanoparticles to deliver fragile antisense oligonucleotides to target the overexpression of PD-L1. We illustrate how the effective knockdown of this receptor can be harnessed to elevate immune cell-mediated killing of aggressive and immunoevasive cancer cells. We observed that MOFs can effectively encapsulate high quantities of various ASO sequences (at *ca.* 9 nmol/mg MOF) while also exhibiting high loading efficiencies. Once encapsulated, ASO@MOF complexes release genetic cargo continually over a 7-day timeframe, which is an advantage that ensures sustained ASO concentrations for longer periods of time and reduces the need for multiple drug administrations in future applications. Upon administration of the treatments to PD-L1-overexpressing tumor cells, MOF encapsulation prevents rapid ASO clearance and degradation and elevates ASO entry into the majority of the cell population. Improved ASO uptake mediated through MOF delivery leads to significant downregulation of overexpressed PD-L1 in two different cancer cells: B16-F10 melanoma and EMT6 triple negative breast cancer. Downregulation of PD-L1 is critically not induced by freely administered ASO treatment or by unloaded MOF, which highlights that the MOF encapsulation of ASO is what promotes their increased potency. Importantly, ASO delivery via MOF nanoparticles has the potential to eliminate the need for: 1) ASO modifications, which can cause detrimental off target binding, require specific tuning for each sequence, and reduce potency, and 2) alternative, commonly used delivery vehicles that often require local administration to target desired cell populations effectively and over extended timeframes.

We observed that ASO@MOF treatment impacted immune cell populations in addition to PD-L1 expression on tumor cells. Namely, ASO@MOF increased immune dendritic cell activation and T cell proliferation. Co-stimulatory marker CD80 and CD86 expression in BMDCs were increased, and CD8^+^ T cells exhibited higher levels of early activation marker CD69 and increased antigen-specific proliferation. Of the three distinct ASO sequences explored in this work, only ASO B was designed in prior literature^14^ to incorporate unique TLR9 agonist modifications within its sequence. However, when delivered herein utilizing MOFs, we observed that all explored ASOs improved immune activation and proliferation. Of note, the measured effects were exhibited at ASO concentrations *ca.* 20-fold lower compared to previous similar studies that employ free ASO.^14^ These changes in immune cell activity propagate when immune cells are co-cultured with ASO@MOF-treated tumor cells, ultimately driving a *ca.* 3-fold greater expression of Caspase-3 compared to immune cells co-cultured with untreated tumor cells. This illustrates that MOF encapsulation of ASOs can both change target gene expression, such as PD-L1, but also enhance innate immune stimulation, which collectively triggers improved antigen-specific anti-tumor immunity. This work highlights how MOF encapsulation can maximize ASO potency on multiple cell types and as such, these findings can be applied to alternative ASO sequences with gene targets beyond PD-L1 and to various cancers or rare diseases. The ability to employ MOFs as carriers for ASO targets will thus positively influence ASO efficacy and broaden the potential for key beneficial immune interactions to overcome aggressive or immunosuppressive diseases.

## MATERIALS AND METHODS

### Materials and Animals

Unless otherwise specified, all materials were purchased commercially and used as received. For ASO synthesis, all materials and reagents were purchased from Glen Research. Water was filtered through a Milli-Q water purification system. Murine B16-F10 (melanoma) and EMT6 (mammary) cell lines were purchased from ATCC and were stored and cultured using recommended conditions. Murine RAW-Blue (macrophage) reporter cells derived from murine RAW 264.7 (macrophage) cells were purchased from InvivoGen and were stored and cultured using recommended conditions. C57BL/6J (wild-type), C57BL/6-Tg (TcraTcrb)1100Mjb/J (OT-1), and B6.Cg-Thy1a/Cy Tg(TcraTcrb)8Rest/J (Pmel-1) mice were purchased from Jackson Laboratory. All national and local guidelines and regulations were followed when handling mice, and all protocols were approved by the Institutional Animal Care and Use Committee (IACUC) at Boston University.

### Cell Culture

B16-F10 cells were cultured according to recommended ATCC conditions with Roswell Park Memorial Institute 1640 Medium (RPMI, Gibco) containing 10% heat-inactivated FBS (HI-FBS, Gibco) and 1% penicillin/streptomycin (Gibco) (termed herein RPMI+/+) in a 37 °C/5% CO_2_ incubator. EMT6 cells were cultured according to recommended ATCC conditions, with Dulbecco’s Modified Eagle Medium (DMEM, ThermoFisher) containing 10% FBS (FBS, Gibco) and 1% penicillin/streptomycin (Gibco) (termed DMEM+/+) in a 37 °C/5% CO2 incubator. RAW-Blue cells were passaged with DMEM+/+ media containing 100 ug/mL Normocin (InvivoGen), and after recovery given 200 ug/mL Zeocin (InvivoGen) every other passage. Cells were passaged and handled according to manufacturer recommendations.

### Synthesis of NU-1000

Stock solutions A (400 mg zirconium chloride) and B (100 mg H4TBAPy linker) were prepared by dissolving each reagent in 50 mL Dimethylformamide (DMF) separately. Both solutions were sonicated for 5 min for full dissolution. 2 mL of stock solution A, 2 mL of stock solution B, 0.8 mL glacial acetic acid and 0.4 mL deionized water were combined in a 2-dram vial. The solution was then capped, briefly vortexed, and immediately placed in a preheated 120 °C oil bath. While in the oil bath, agitation of the yellow solution was avoided. When the solution began to turn cloudy, after c.a. 3-5 minutes, it was removed and put into an ice bath. After cooling to room temperature (RT), NU-1000 MOF nanoparticles were isolated via centrifugation at 8000 RPM for 20 min, and then washed with 5 mL DMF. Each reaction batch yielded c.a. 4 mg of NU-1000, which was minimally variable and relatively homogenous, and imaged using scanning electron microscopy (SEM) to determine batch particle size (Fig. S1). Batches with similar particle sizes were combined (within a range of 100 – 200 nm) and washed twice with DMF with a 1 h soaking period between washes. After the second wash, combined batches were washed with an acidic DMF solution (1.2 mL DMF and 0.05 mL 8M aq. HCl per batch) and heated at 100 °C for 18 h. Once at RT, the MOF was washed 3 times with DMF (4 mL per batch), and then acetone (4 mL per batch), with a 1 h soaking period between washes. For long term storage, NU-1000 was kept in acetone. Overnight thermal activation in a vacuum oven at 80 °C was done immediately before MOF use.

### Synthesis and Purification of ASOs

ASOs were synthesized using a MerMade 12 oligonucleotide synthesizer (LGC Biosearch Technologies) on controlled pore glass beads (Universal UnyLinker Support 1000Å) using standard phosphoramidite chemistry and a phosphorothioate backbone. Fluorophore-labelled ASO A was tagged with Cy3 dye (Cyanine 3 Phosphoramidite, Glen Research) via a hand-coupling procedure. Cy3 dye hand-coupling was performed by first washing ASOs removed from the MerMade with 10 mL Deblock (3% trichloroacetic acid in dichloromethane) followed by 20 mL anhydrous acetonitrile (ACN), then coupling the Cy3 dye with activator (0.25M 5-ethylthio-1H-tetrazole (ETT) in anhydrous acetonitrile) for 15 min while protected from light. Cy3 dye-labelled ASO was then washed with 20 mL ACN, sulfurized with 10 mL sulfurizing reagent (0.05M Sulfurizing Reagent II in Pyridine/Acetonitrile) for ∼30 sec, and washed a second time with 20 mL ACN. Sequence was then capped with 2 mL of Cap A (Tetrahydrofuran/acetic anhydride/2,6-lutidine (80/10/10)) and Cap B (1-Methylimidazole/tetrahydrofuran (16/84)), washed with 20 mL ACN, dried by blowing air through the column and returned to the MerMade synthesizer to complete final base pair additions. Post-synthesis, strands were deprotected using 3 mL of a 1:1 solution of 37% ammonium hydroxide:40% methylamine (Sigma) at 55 °C for 30 min, or with 3 mL of 37% ammonium hydroxide overnight at room temperature (RT) for dye-labeled strands. Strands were purified via reverse-phase high-performance liquid chromatography (RP-HPLC, Agilent) using a gradient of 0.1 M triethylammonium acetate and 3% ACN in water to 100% ACN. Purification was performed using either a C18, or C3 if dye-labeled, column (Agilent). Fractioned samples were incubated in a 20% aqueous acetic acid solution at RT for 1 h, then washed 3 times with ethyl acetate (Sigma) to remove the protective dimethoxytrityl groups. DNA products were lyophilized and resuspended in water. Matrix-assisted laser desorption/ionization time-of-flight (MALDI-TOF; Bruker) mass spectrometry (matrix: 2’,4’, -dihydroxyacetophenone) was used to verify the DNA products by comparing the molecular weight of the analyte to IDT OligoAnalyzer Tool calculations. Ultraviolet-visible (UV-Vis, Agilent Cary 60 spectrophotometer) absorption was measured at 260 nm, or 555 nm for dye-labeled strands, to quantify ASO concentrations.

### ASO Encapsulation into NU-1000

ASOs were prepared at 1 μM in 100 μL water. 0.1-0.2 mg of NU-1000 MOF per encapsulation was measured, resuspended in 500 μL water, and sonicated prior to centrifugation at 14800 RPM for 1 min to wash. Supernatant liquid was removed from the pelleted MOF sample and discarded. 100 μL of the ASO solution was added to the MOF pellet, and the sample was vortexed, sonicated, and centrifuged at 14800 RPM for 1 min. The sample was placed on a heat block at 37°C and 300 RPM for 3.5 h. After incubation, the sample was centrifuged for 1 min at 14800 RPM. Supernatant was removed and was used to quantify amount ASO loaded, where it was stored at −20 °C prior to measurement. Encapsulated samples were stored at 4 °C until use. Supernatant ASO concentration was measured through UV-Vis absorption at 260 nm, or 555 nm for dye-labeled strands, using the extinction coefficients calculated by the IDT OligoAnalyzer Tool. To determine encapsulation efficiency, the difference between the number of moles of ASO in the initial incubation solution and the final supernatant solution was calculated.

### NU-1000 ASO Release Studies

0.2 mg of Cy3 dye-labelled ASO A@MOF was incubated in 1 mL of 1x PBS in a heat block at 37°C and 300 RPM. At specified timepoints, the MOF sample was centrifuged at 14,800 RPM for 1 min to collect and replace supernatant with 1 mL PBS for analysis. Supernatant was stored at −80°C. Once all timepoint samples were collected, supernatants were analyzed using UV-Vis at 555 nm to quantify the release over time. Percent release was determined by normalizing based on the initial amount of ASO loaded.

### ASO@MOF Cellular Uptake

B16-F10 cells were seeded at 5×10^4^ cells/well in a 48-well flat bottom plate and allowed to adhere overnight. Cells were washed with 1x PBS and treated with 250 μL of the different conditions at 1 μM in media. Treatments of MOF, dye-labelled ASO A, and dye-labelled ASO A@MOF were prepared immediately before use by resuspending in 1 mL of RPMI+/+. Cells were treated for 24 h and then washed with 1x PBS. Cells were detached using 1x TrypLE™ Express Enzyme (ThermoFisher) and transferred to microtiter tubes using 600 μL of 1x PBS to wash. Cells were centrifuged at 1200 RPM for 5 min and washed with 1x PBS. Cells were then stained with Fixable Live/Dead-Far Red (Invitrogen) for 20 min at 4 °C, after which they were washed with 1x PBS, fixed using 100 μL 4% paraformaldehyde fixation buffer (BioLegend), and incubated at 4 °C until measurement with an Attune NxT flow cytometer. Uptake was analyzed by Cy3 fluorescence for ASO and by inherent MOF fluorescence at 387 nm from a 488 nm excitation laser. Data were analyzed via FlowJo.

### *In vitro* PD-L1 Surface Expression Knockdown

B16-F10 cells were seeded at 5×10^4^ cells/well in a 48-well flat bottom plate and allowed to adhere overnight. Cell media was replaced with RPMI+/+ containing 150 ng/mL of recombinant mouse interferon gamma (IFNγ, BioLegend) to stimulate PD-L1 overexpression. After a 24 h incubation, the media was replaced with 250 μL RPMI+/+ containing MOF, ASO, or ASO@MOF prepared immediately before administration at 1 μM concentrations. EMT6 cells were prepared and treated similarly. After treatment for 24 h, cells were washed and transferred to microtiter tubes as previously described in the ASO@MOF Cellular Uptake section. Cells were stained with Fixable Live/Dead-Far Red (Invitrogen) and PE anti-mouse PD-L1 (BioLegend, #155404), before washing and resuspending in 100 μL 4% paraformaldehyde fixation buffer as previously described in the ASO@MOF Cellular Uptake section. Flow cytometry analysis was performed with an Attune NxT flow cytometer and data were analyzed via FlowJo.

### BMDC Collection

Hindlegs of C57BL/6 mice were harvested for bone marrow-derived dendritic cell (BMDC) collection. Using a needle and syringe of RPMI+/+, cells were flushed from bone marrow into a 100 mm petri dish. The cell suspension was centrifuged at 1200 RPM for 5 min and supernatant was aspirated. Red blood cells were lysed using 2 mL of ACK lysing buffer (Gibco) for 4 min at RT. Cells were washed with 1x PBS and cultured in 100 mm petri dishes (Nunclon) in 20 mL of RPMI+/+ with 40 ng/mL granulocyte-macrophage colony-stimulating factor (GM-CSF, BioLegend) for 7 days in an incubator. 5 mL of RPMI+/+ was added after day 4 to maintain nutrients.

### *In vitro* Co-Stimulatory Surface Marker Expression

7 days after BMDC collection, cells were scraped and transferred to a 96-well round bottom plate at 0.2×10^6^ cells/well and allowed to recover for 1 h. MOF, ASO, and ASO@MOF treatments were prepared in RPMI+/+ media. After cell recovery, 1 μM treatments were added at 100 μL for a total volume of 200 μL in each well, and cells were returned to the incubator. After 24 h, cells were transferred to microtiter tubes and washed once with 1x PBS. Cells were centrifuged at 1200 RPM for 5 min. Supernatant was removed and cells were stained with Fixable Live/Dead-Far Red (Invitrogen), CD11c—PE (BioLegend, #117308), PerCP/Cyanine5.5 anti-mouse CD80 (BioLegend, #104722), and PE/Cyanine7 anti-mouse CD86 (BioLegend, #105014) for 20 min at 4 °C. Cells were washed again with 1x PBS, resuspended in 100 μL 4% paraformaldehyde fixation buffer, and incubated at 4 °C until flow cytometry analysis. Co-stimulatory marker expression was determined in CD11c^+^ BMDCs based on fluorescence intensity of the antibodies.

### *In vitro* Splenocyte Proliferation and Activation

BMDCs were obtained as previously described, scraped, and transferred at 3×10^4^ cells/well in a 96-well round bottom plate. After 1 h recovery in an incubator, cells were treated with unloaded MOF, free ASO, or ASO@MOF at a 1 μM concentration in a 50 μL volume. After 24 h, 50 μL volume of 1 μM treatments of unloaded MOF, free ASO, or ASO@MOF were again added to cells with an additional 50 μL of 100 μg/mL of Endofit Ovalbumin (Invivogen). After another 24 h, a spleen was harvested from an OT-I mouse, and a single cell suspension was obtained by straining through a 70 μM strainer with continuous PBS flow. Cells were centrifuged at 1200 RPM for 5 min and supernatant was aspirated. To selectively lyse red blood cells, 3 mL of ACK lysing buffer was used at RT for 5 min. Cells were then washed with 1x PBS, centrifuged at 1200 RPM for 5 min, and resuspended at 1 ×10^8^ cells/mL in 1x MojoSort Buffer (BioLegend). CD8^+^ T cells were isolated using an EasySep Mouse CD8^+^ T Cell Isolation Kit and the manufacturer’s protocol (STEMCELL Technologies). Isolated CD8^+^ T cells were resuspended in 500 μL 1x PBS and stained with 500 μL of eFluor 670 (Invitrogen) following the manufacturer’s protocol. After staining, cells were resuspended in RPMI+/+ at 3×10^6^ cells/mL. The 96-well plate containing BMDCs was centrifuged at 1200 RPM for 5 min and 100 μL of supernatant was removed. Efluor 670-stained CD8^+^ T cells were added to BMDC cells at a final ratio of 1:5 BMDCs to CD8^+^ T cells, which were co-cultured for 72 h. Cells were then transferred to microtiter tubes, washed with 1x PBS, stained with fluorescent-conjugated antibodies PE/Cyanine7 anti-mouse CD69 (BioLegend, # 104511) and PE anti-mouse CD8a (BioLegend, # 162303), and analyzed with live flow cytometry. Histograms of eFluor 670 emission were used to measure the percentage of proliferating CD8^+^ T cells with a negative control gate at *ca.* <5% proliferation of an untreated group. Early activation of CD8^+^ T cells was quantified through percent of CD69 positive CD8^+^ T cells.

### *In vitro* Analysis of ASO Activation on Immune Cells

RAW-blue cells were passaged and resuspended in DMEM+/+ media without any additional antibiotics at 2×10^6^ cells in a T-75 culture flask 3 days before experimental use. After 3 days, media was removed, cells were washed twice with 1x PBS, detached with a cell scraper, and resuspended at 5.5×10^5^ cells/mL in DMEM with 10% heat-inactivated fetal bovine serum. Cells were seeded in a 96-well flat bottom plate at 180 μL cell stock and 20 μL treatment. Treatments of ASOs and positive control CpG ODN 1826 were prepared immediately before use in DMEM with 10% heat inactivated fetal bovine serum. After overnight incubation of cells, QUANTI-Blue Solution (InvivoGen) was prepared according to the manufacturer instructions. In a separate 96-well flat bottom plate, 180 μL of Quanti-Blue was combined with 20 μL of cell supernatant, with a media only background control. Quanti-Blue assay was incubated at 37 °C for 1 h, and then optical density was read at 620 nm using a SpectraMax i3X plate reader (Molecular Devices). A dose response curve was plotted to identify concentration-dependent activation.

### *In vitro* Active Caspase-3 Analysis to Quantify Tumor Cell Death

B16-F10 cells were passaged and resuspended in RPMI +/+ media containing 150 ng/mL IFNγ for 24 h, then collected and stained with eFluor 670 (Invivogen) as previously described in the *In vitro* Splenocyte Proliferation and Activation section. Cells were then seeded in a 96-well flat bottom plate at 8×10^3^ cells/well and treated with ASO, ASO@MOF or unloaded MOF at 1 μM concentrations for 24 h. Pmel-1 splenocytes were harvested as previously described in the *In vitro* Splenocyte Proliferation and Activation section by obtaining a single cell suspension from a harvested Pmel-1 spleen, selectively lysing red blood cells with ACK buffer, and then resuspending the remaining splenocyte population at 4×10^6^ cells/mL. Media containing Pmel-1 splenocytes was used to replace the treatment media of B16-F10 cells for a final ratio of 100:1 effector Pmel-1 splenocytes to target B16-F10 tumor cells at 200 μL total volume/well. Cells were co-cultured for 50 h and then stained with PE Active Caspase-3 anti-mouse (Fisher Scientific, #BDB570183) using BD Cytofix/Cytoperm™ Fixation/Permeabilization Kit (BD Biosciences, #554714) following manufacturer instructions. Flow cytometry analysis was performed to determine the extent of Active Caspase-3 in tumor cell populations.

### Statistical Analysis

All values shown in graphs are mean ± standard deviation. Statistical analysis was performed using GraphPad Prism 9 software, with experiment sample sizes described in figure captions. Comparisons between two groups used unpaired t-tests, and comparisons between more than two groups used an ordinary one-way ANOVA with an appropriate post-hoc multiple comparison test. No specific pre-processing of data was performed prior to analyses. Significance was defined as P < 0.05 (*P < 0.05; **P < 0.01; ***P < 0.001; ****P < 0.0001; ns = not significant).

## Acknowledgements

The authors would like to thank Dr. Xin Brown for training on core equipment, and Dr. Yijing Chen for the initial development of nano NU-1000 synthesis. M.H.T. acknowledges financial support from Boston University through startup funding support. This work was supported in part by the Arnold and Mabel Beckman Foundation through a Beckman Young Investigator Award, and the National Institute of General Medical Sciences of the National Institutes of Health award R35GM157326. J.A.N. acknowledges financial support from the Boston University Biological Design Center, STEM Pathways Program (DoD STEM FY20 Award HQ00342110008), and Distinguished Summer Research Fellowship. J.A.N. and L.B acknowledge financial support from the Boston University Undergraduate Research Opportunities Program. E.C. acknowledges financial support from an NSF GRFP and Translational Research in Biomaterials T32 training grant (T32EB006359). M.A.D. acknowledges financial support from ACS IRG 22-153-42 from the American Cancer Society. S.Z. acknowledges financial support from The Hartwell Foundation and Synthetic Biology and Biotechnology (SB2) Predoctoral Training Grant (T32GM130546). F.S., J.S.M, and O.K.F. acknowledge financial support of the CBC-NU Cell-free Biomanufacturing Institute funded by the U.S. Army Contracting Command Award W52P1J-21-9-3023. F.S. also gratefully acknowledges support from the Ryan Fellowship and the International Institute for Nanotechnology at Northwestern University.

## REFERENCES

1. Yamaguchi, H., Hsu, J.-M., Yang, W.-H. & Hung, M.-C. Mechanisms regulating PD-L1 expression in cancers and associated opportunities for novel small-molecule therapeutics. Nat. Rev. Clin. Oncol. 19, 287–305 (2022).

2. Iwai, Y. et al. Involvement of PD-L1 on tumor cells in the escape from host immune system and tumor immunotherapy by PD-L1 blockade. Proc. Natl. Acad. Sci. 99, 12293–12297 (2002).

3. Ju, X., Zhang, H., Zhou, Z. & Wang, Q. Regulation of PD-L1 expression in cancer and clinical implications in immunotherapy. Am. J. Cancer Res. 10, 1–11 (2020).

4. Zerdes, I., Matikas, A., Bergh, J., Rassidakis, G. Z. & Foukakis, T. Genetic, transcriptional and post-translational regulation of the programmed death protein ligand 1 in cancer: biology and clinical correlations. Oncogene 37, 4639–4661 (2018).

5. Jiang, Y., Chen, M., Nie, H. & Yuan, Y. PD-1 and PD-L1 in cancer immunotherapy: clinical implications and future considerations. Hum. Vaccines Immunother. 15, 1111–1122 (2019).

6. Beck, A., Wurch, T., Bailly, C. & Corvaia, N. Strategies and challenges for the next generation of therapeutic antibodies. Nat. Rev. Immunol. 10, 345–352 (2010).

7. Sharma, P., Hu-Lieskovan, S., Wargo, J. A. & Ribas, A. Primary, Adaptive, and Acquired Resistance to Cancer Immunotherapy. Cell 168, 707–723 (2017).

8. Nishino, M., Ramaiya, N. H., Hatabu, H. & Hodi, F. S. Monitoring immune-checkpoint blockade: response evaluation and biomarker development. Nat. Rev. Clin. Oncol. 14, 655– 668 (2017).

9. Topalian, S. L. et al. Safety, Activity, and Immune Correlates of Anti–PD-1 Antibody in Cancer. N. Engl. J. Med. 366, 2443–2454 (2012).

10. Castiglioni, A. et al. Combined PD-L1/TGFβ blockade allows expansion and differentiation of stem cell-like CD8 T cells in immune excluded tumors. Nat. Commun. 14, 4703 (2023).

11. Roberts, T. C., Langer, R. & Wood, M. J. A. Advances in oligonucleotide drug delivery. Nat. Rev. Drug Discov. 19, 673–694 (2020).

12. Gagliardi, M. & Ashizawa, A. T. The Challenges and Strategies of Antisense Oligonucleotide Drug Delivery. Biomedicines 9, 433 (2021).

13. Chou, L., Callmann, C. E., Dominguez, D., Zhang, B. & Mirkin, C. A. Disrupting the Interplay between Programmed Cell Death Protein 1 and Programmed Death Ligand 1 with Spherical Nucleic Acids in Treating Cancer. ACS Cent. Sci. 8, 1299–1305 (2022).

14. Fernandez-Rodriguez, L. et al. Dual TLR9 and PD-L1 targeting unleashes dendritic cells to induce durable antitumor immunity. J. Immunother. Cancer 11, e006714 (2023).

15. Zhu, Y., Zhu, L., Wang, X. & Jin, H. RNA-based therapeutics: an overview and prospectus. Cell Death Dis. 13, 1–15 (2022).

16. Liang, X.-H., Sun, H., Nichols, J. G. & Crooke, S. T. RNase H1-Dependent Antisense Oligonucleotides Are Robustly Active in Directing RNA Cleavage in Both the Cytoplasm and the Nucleus. Mol. Ther. J. Am. Soc. Gene Ther. 25, 2075–2092 (2017).

17. Crooke, S. T., Baker, B. F., Crooke, R. M. & Liang, X. Antisense technology: an overview and prospectus. Nat. Rev. Drug Discov. 20, 427–453 (2021).

18. Huang, S. et al. Nonviral delivery systems for antisense oligonucleotide therapeutics. Biomater. Res. 26, 49 (2022).

19. Deleavey, G. F. & Damha, M. J. Designing Chemically Modified Oligonucleotides for Targeted Gene Silencing. Chem. Biol. 19, 937–954 (2012).

20. Lauffer, M. C., van Roon-Mom, W. & Aartsma-Rus, A. Possibilities and limitations of antisense oligonucleotide therapies for the treatment of monogenic disorders. Commun. Med. 4, 1–11 (2024).

21. Anderson, B. A. et al. Towards next generation antisense oligonucleotides: mesylphosphoramidate modification improves therapeutic index and duration of effect of gapmer antisense oligonucleotides. Nucleic Acids Res. 49, 9026–9041 (2021).

22. Tanaka, H. et al. Delivery of Oligonucleotides Using a Self-Degradable Lipid-Like Material. Pharmaceutics 13, 544 (2021).

23. Gao, P., et al. Antitumor Agents Based on Metal–Organic Frameworks. Angew. Chem. Int. Ed. 60, 16763–16776 (2021).

24. He, S. et al. Metal-organic frameworks for advanced drug delivery. Acta Pharm. Sin. B 11, 2362–2395 (2021).

25. Sun, Y. et al. Metal–Organic Framework Nanocarriers for Drug Delivery in Biomedical Applications. Nano-Micro Lett. 12, 103 (2020).

26. Tong, P.-H. et al. Metal–organic frameworks (MOFs) as host materials for the enhanced delivery of biomacromolecular therapeutics. Chem. Commun. 57, 12098–12110 (2021).

27. Teplensky, M. H. et al. Temperature Treatment of Highly Porous Zirconium-Containing Metal–Organic Frameworks Extends Drug Delivery Release. J. Am. Chem. Soc. 139, 7522– 7532 (2017).

28. Abazari, R. et al. Design and Advanced Manufacturing of NU-1000 Metal–Organic Frameworks with Future Perspectives for Environmental and Renewable Energy Applications. Small 20, 2306353 (2024).

29. Ni, K., Luo, T., Nash, G. T. & Lin, W. Nanoscale Metal–Organic Frameworks for Cancer Immunotherapy. Acc. Chem. Res. 53, 1739–1748 (2020).

30. Della Rocca, J., Liu, D. & Lin, W. Nanoscale Metal–Organic Frameworks for Biomedical Imaging and Drug Delivery. Acc. Chem. Res. 44, 957–968 (2011).

31. Concha-Benavente, F. et al. Identification of the cell-intrinsic and extrinsic pathways downstream of EGFR and IFNγ that induce PD-L1 expression in head and neck cancer. Cancer Res. 76, 1031–1043 (2016).

32. Wang, T. C. et al. Scalable synthesis and post-modification of a mesoporous metal-organic framework called NU-1000. Nat. Protoc. 11, 149–162 (2016).

33. Horcajada, P. et al. Metal–Organic Frameworks in Biomedicine. Chem. Rev. 112, 1232– 1268 (2012).

34. Teplensky, M. H. et al. A Highly Porous Metal-Organic Framework System to Deliver Payloads for Gene Knockdown. Chem 5, 2926–2941 (2019).

35. Peng, Q. et al. PD-L1 on dendritic cells attenuates T cell activation and regulates response to immune checkpoint blockade. Nat. Commun. 11, 4835 (2020).

