## Supplemental Information for "Strengthening Antisense Oligonucleotide-Mediated Anti-Tumor Immunity via Metal-Organic Framework Nanoparticles"

**This PDF file includes:**

Figures S1 to S4
Table S1

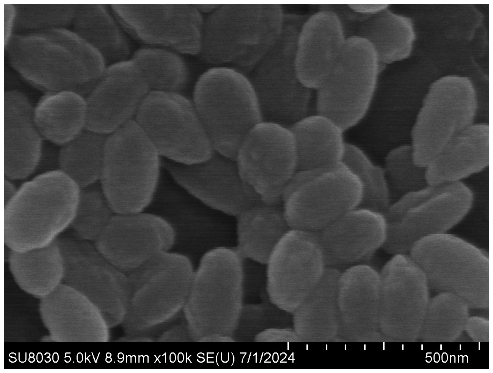


**Figure S1.** Representative scanning electron microscope (SEM) image of nanosized NU-1000.

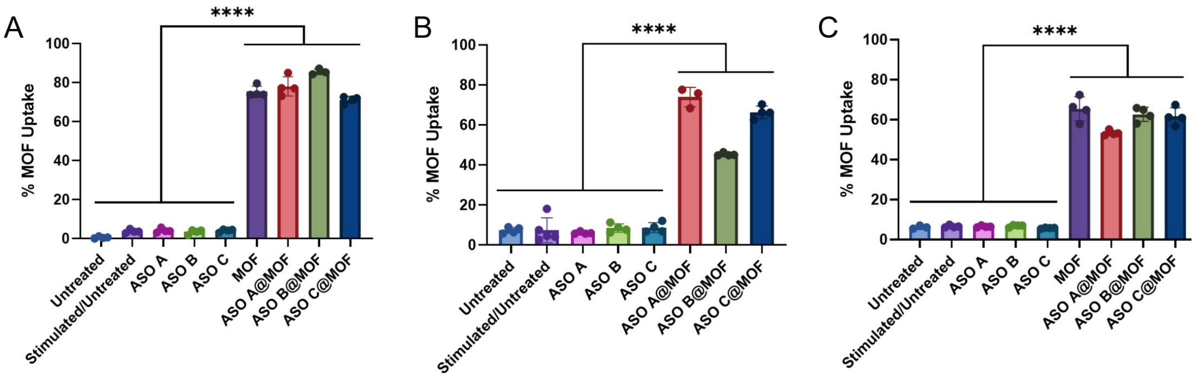


**Figure S2.** MOF entry into cells is cell type independent. Statistically equivalent cellular uptake for MOFs encapsulating various ASO cargo occurs in **(A)** B16-F10 melanoma, **(B)** EMT6 triple negative breast cancer, and **(C)** bone marrow-derived dendritic cells.

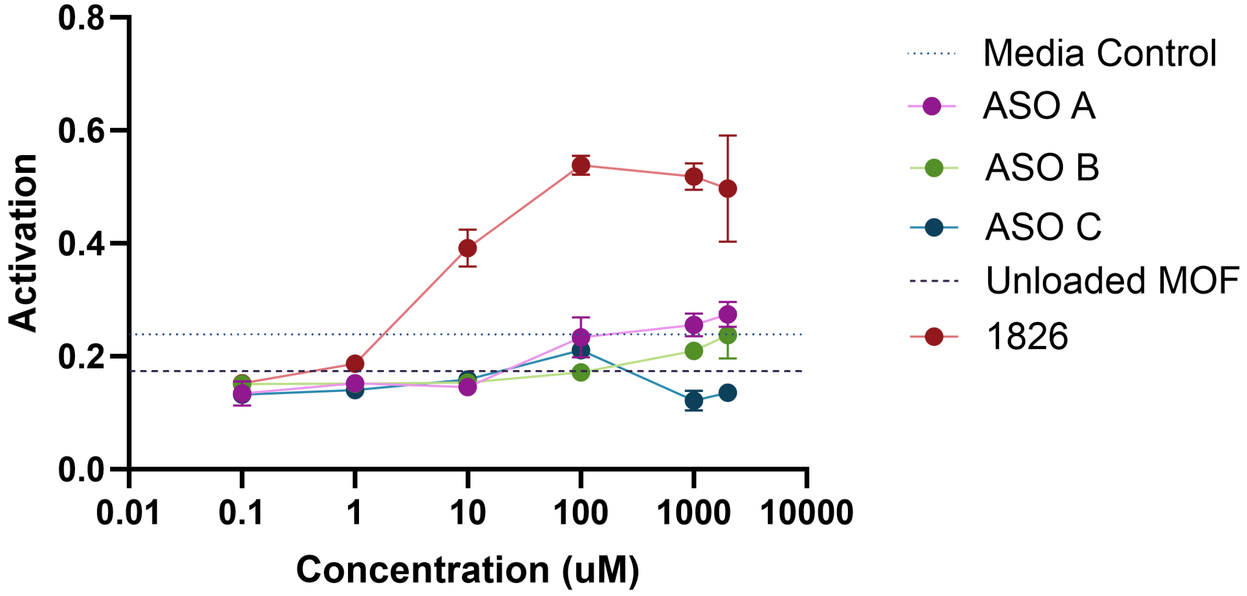


**Figure S3.** Activation of RAW Blue Macrophage reporter cells with various ASO sequences, unloaded NU-1000 MOF control, or positive control CpG 1826 agonist.

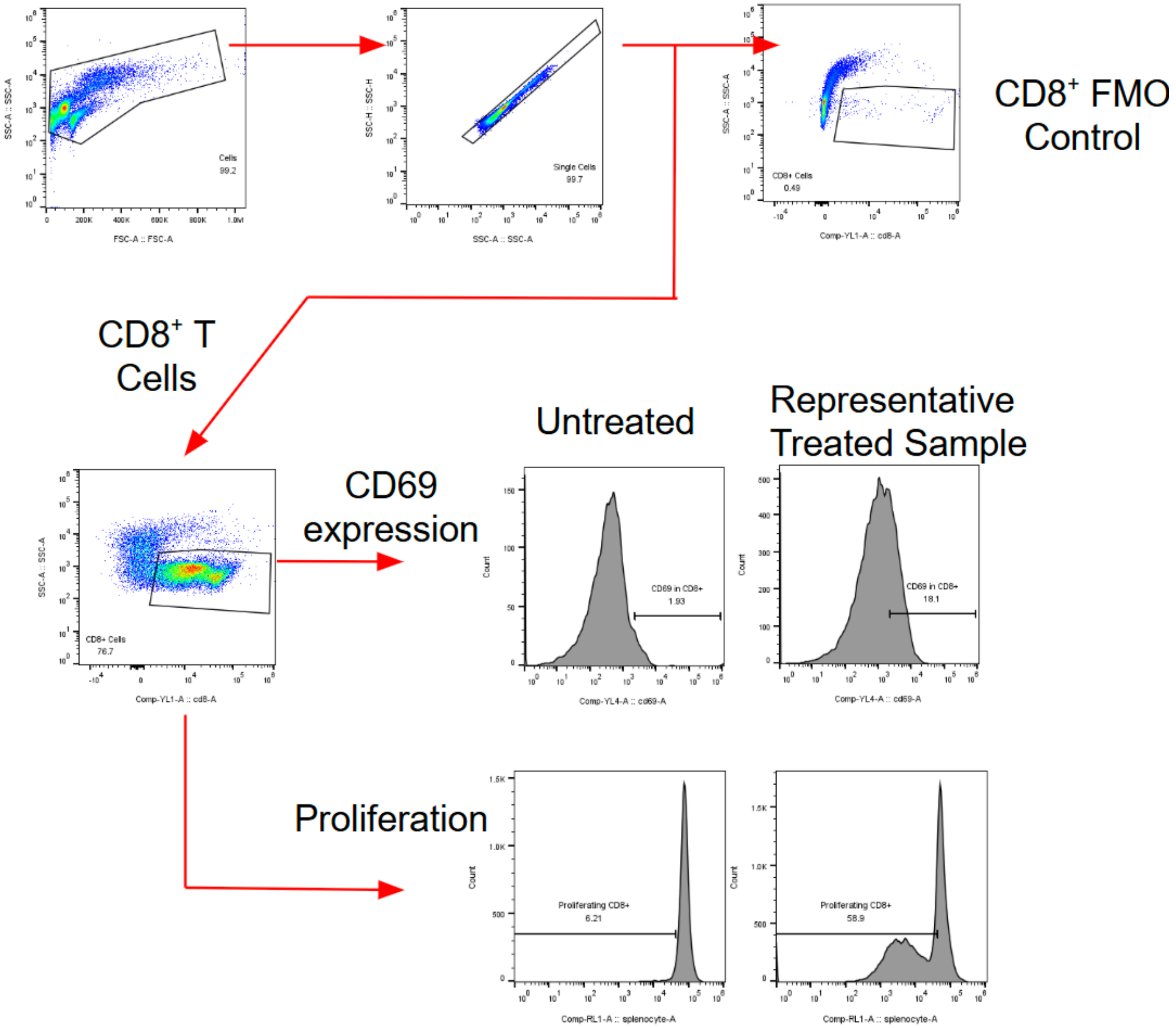


**Figure S4.** Gating strategy to identify proliferation and activation of CD8$^+$ T cells. Splenocytes were gated via fluorescent antibody staining to identify CD8$^+$ T cells and their CD69 expression, and via proliferation dye dilution to identify T cell proliferation.

**Table S1.** Summary information of ASO sequences used in this work.

| Strand Name | Sequence (5' to 3')[1] | Expected Mass (g/mol)[2] | Experimental Mass (g/mol) (measured via MALDI-TOF) |
|---|---|---|---|
| ASO A | TGA CGT TGC TGC CAT | 4800 | 4808 |
| ASO B | CTT ACG TCT CCT CGA | 4719.9 | 4788 |
| ASO C | GTT GAT TTT GCG GTA T | 5190.3 | 5235 |
| Dye-labelled ASO A | TGA CGT TGC TGC CAT (Cy3)[3] TT | 5963 | 5946.3 |

[1]All sequences utilized a phosphorothioate backbone
[2]Calculated using IDT's OligoAnalyzer Tool: https://www.idtdna.com/calc/analyzer
[3]1-[3-(4-monomethoxytrityloxy)propyl]-1'-[3-[(2-cyanoethyl)-(N,N-diisopropyl)phosphoramidityl]propyl]-3,3,3',3'-tetramethylindocarbocyanine chloride (Glen Research #182873-76-3)
